# Have It, Know It, but Don’t Show It: Examining Physiological Arousal, Anxiety, and Facial Expressions over the Course of a Social Skills Intervention for Autistic Adolescents

**DOI:** 10.1101/582676

**Authors:** Niharika Jain, Sheikh Iqbal Ahamed, Serdar Bozdag, Bridget K. Dolan, Alana J. McVey, Kirsten S. Willar, Sheryl S. Pleiss, Christina C. Murphy, Christina L. Casnar, Stephanie Potts, Daniel Cibich, Kylie Nelsen-Freund, Dana Fernandez, Illeana Hernandez, Amy Vaughan Van Hecke

## Abstract

Facial expressions provide a nonverbal mechanism for social communication, a core challenge for autistic people. Little is known regarding the association between arousal, self-report of anxiety, and facial expressions among autistic adolescents. Therefore, this study investigated session-by-session facial expressions, self-report of anxiety, and physiological arousal *via* Electrodermal Activity (EDA), of 12 autistic male adolescents in a didactic social skills intervention setting. The goals of this study were threefold: 1) identify physiological arousal levels (“have-it”), 2) examine if autistic adolescents’ facial expressions indicated arousal (“show-it”), and 3) determine whether autistic adolescents were self-aware of their anxiety (“know-it”). Our results showed that autistic adolescents’ self-rated anxiety was significantly associated with peaks in EDA. Both machine learning algorithms and human participant-based methods, however, had low accuracy in predicting autistic adolescents’ arousal state from facial expressions, suggesting that autistic adolescent’s facial expressions did not coincide with their arousal. Implications for understanding social communication difficulties among autistic adolescents, as well as future targets for intervention, are discussed. This project is registered with ClinicalTrials.gov, Identifier: NCT02680015.

---

Autism spectrum disorder is a lifelong developmental condition characterized by challenges in social interaction and communication frequently coinciding with the presence of restricted and/or repetitive behaviors (American Psychiatric Association, 2013). Heterogeneity in symptom presentation of people with autism spectrum disorder, hereafter referred to as autistic people^*^ makes diagnosis, treatment, and intervention more challenging (American Psychiatric Association, 2013). In addition, anxiety is considered to be one of the most common co-occurring difficulties among autistic people (van Steensel, Bögels & Perrin, 2011; White, Oswald, Ollendick, & Scahill, 2009; White et al., 2010). Several studies suggest that autistic children, versus typically developing (TD) children, are at an increased risk of developing anxiety disorders (Kim, Szatmari, Bryson, Streiner, & Wilson, 2000; Kuusikko et al., 2008; Leyfer et al., 2006; Mayes, Calhoun, Murray, Ahuja, & Smith, 2011). The presence of anxiety among autistic youth may lead to more complex behaviors such as aggressive and oppositional behaviors (Kim et al., 2000), exacerbated challenges with social functioning (de Bruin, Ferdinand, Meester, de Nijs, & Verheij, 2007), and more distressing thoughts (Farrugia & Hudson, 2006). If these cognitive and behavioral challenges are not addressed in youth, they may become persistent and more difficult to treat, leading to poorer overall outcomes in adulthood (Magiati, Tay, & Howlin, 2014). The assessment of anxiety disorders among autistic people is complex, because of the overlapping of symptomatology between anxiety and autism, inherent communication difficulties in autism, and the idiosyncratic nature of these disorders (Davis, Saeed, & Antonacci, 2008; White et al., 2009). This may be further complicated by high rates of alexithymia, a difficulty describing one’s own emotional state, in autism; approximately half of autistic people have been shown to experience alexithymia (Milosavljevic et al., 2016; Paula-Pérez, Martos-Pérez, & Llorente-Comí, 2010; Shah, Hall, Catmur, & Bird, 2016).

In the present study, a brief review of the physiological and behavioral measurement of anxiety and arousal in autism is offered. These findings are taken in light of the current study, which examined session-by-session physiological arousal using electrodermal activity (EDA), self-rated anxiety symptoms, and facial expressions among a group of autistic adolescents participating in a social skills intervention. The relation among the aforementioned factors was studied to see if the adolescents: 1) experienced physiological arousal in a naturalistic environment, 2) were self-aware of their anxiety, and 3) demonstrated anxiety *via* their facial expressions. Finally, comparison of image classification accuracy by machine learning methods and human participants was conducted.

Anxiety is known to have both cognitive and somatic components (Liebert & Morris, 1967). Cognitive, or mental, components are associated with the expectation of “negative” events and, consequently, unsuccessful performance (Vickers & Williams, 2007; Vitasari et al., 2011). Somatic components, on the other hand, are associated with the arousal of the autonomic nervous system (ANS) or physiological arousal (Vickers & Williams, 2007; Vitasari et al., 2011). There are a variety of approaches for studying physiological arousal and coinciding emotional processes. A review by (White et al., 2014) provides a comprehensive evaluation of such approaches, listing startle responses, EDA (also referred to as skin conductance response; SCR), and cardiac activity as some of the most common techniques employed for understanding emotion regulation difficulties pertaining to anxiety in autism. Electrodermal activity serves as a reliable measure of sympathetic nervous system (SNS) activity (Critchley, 2009), which may be helpful in understanding the arousal state of a person, especially among those that may struggle to otherwise communicate their own arousal.

Although there has been considerable work done examining EDA as a measure of arousal among autistic youth (see Table 1), the results of these studies are not consistent and, hence, do not provide a clear understanding of the role played by EDA in emotion regulation of autistic youth. A possible explanation for the inconsistent findings is the variability in the paradigms used. Only two (out of 14; Levine et al., 2012; South, Dana, White, & Crowley, 2011) studies utilized a stressor task to elicit emotions, while the remaining 12 studies employed photographs for emotional arousal. Moreover, all of these studies compared the EDA levels of autistic people to those of TD control groups. No known study, to date, has examined EDA data as a base to understand the emotions and arousal experienced by autistic adolescents in a naturalistic setting nor longitudinally. Both of these types of investigations are needed, as laboratory studies of EDA in autism in response to stimuli may not provide an adequate conceptualization of the day-to-day functioning of the person. More specifically, an unfamiliar, one-time laboratory measurement of EDA may already be confounded by the fear or anxiety that novel situations elicit in autistic people (Gillott, Furniss, & Walter, 2001).

**Table 1.**
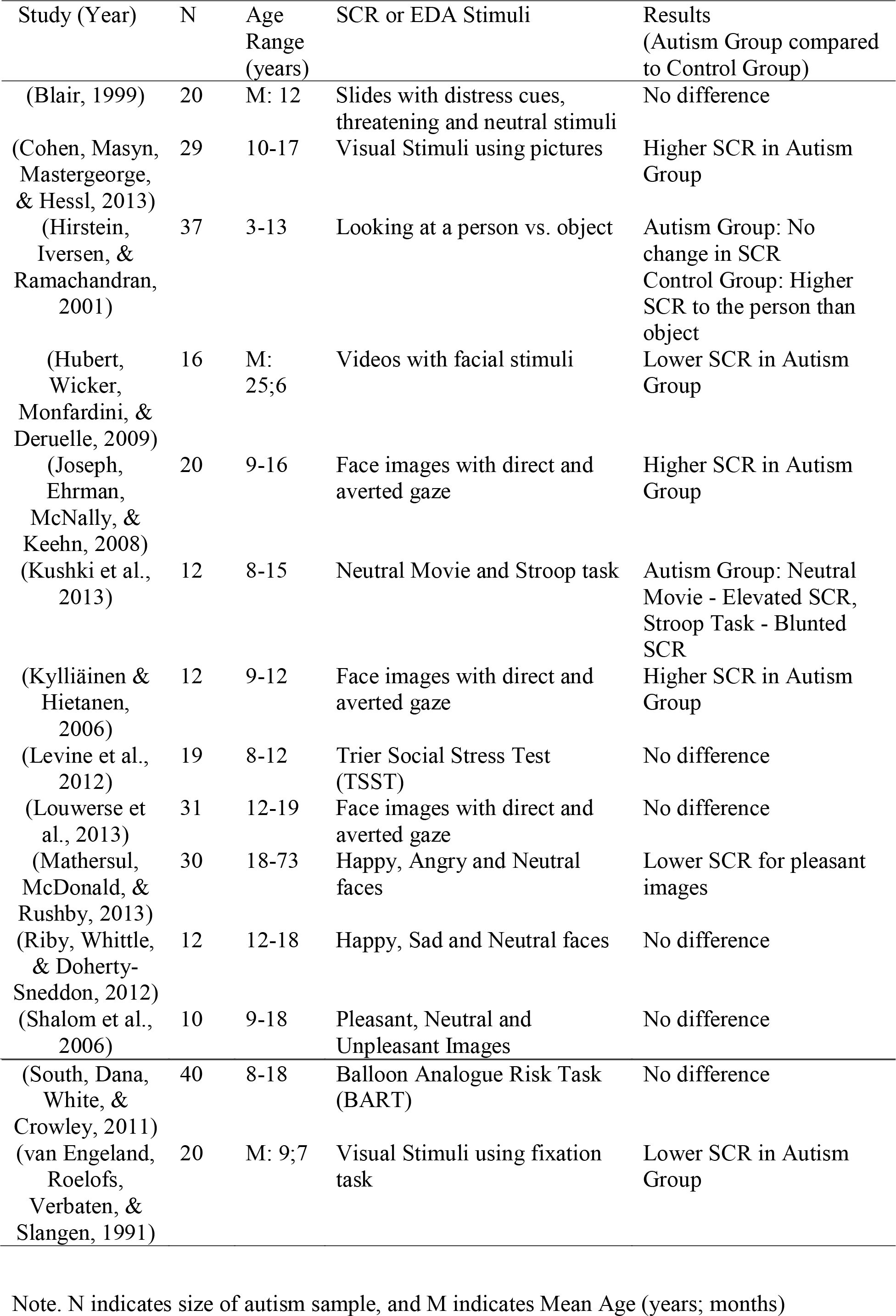
Studies of Skin Conductance Responses among Autistic Individuals

Beyond SNS activity, there are behavioral methods of assessing the arousal of another person, namely, through facial expressions. Facial expressions have long been studied as a means of understanding emotions (Ekman, 1993). The time-consuming and laborious micro-coding of facial expressions among autistic youth, however, has perhaps contributed to the limited research in this area. New methods have attempted to develop automated, computerized systems for facial expression identification. The automated analysis of facial expressions is a complex process that consists of 1) detection of the face, 2) extraction of facial features, and 3) classification of facial expressions (Pantic & Rothkrantz, 2000). The Facial Action Coding System (FACS) (Ekman & Friesen, 1978) is the most commonly used technique to study facial activity, but FACS has its own set of limitations, irrespective of the classification method used, manual or automated (Pantic & Rothkrantz, 2000). Combinations of several techniques, Parameterized Appearance Modeling (De la Torre & Cohn, 2011), Eigenfaces (Turk & Pentland, 1991), Principal Component Analysis (Hotelling, 1933), classification methods (Pantic & Rothkrantz, 2000), and clustering (Jain & Dubes, 1988), may be employed for a more effective analysis of facial expressions (De la Torre & Cohn, 2011; Pantic & Rothkrantz, 2000).

As in the case of the existing EDA studies described above, the literature pertaining to emotion perception and the production of facial expressions by autistic people fails to provide a consensus (Kennedy & Adolphs, 2012). There are two established approaches for emotion-based studies. One approach is to determine the emotion perception or emotion recognition of autistic people. The second approach is to examine emotion experiences, *via* facial expressions, of autistic people. Researchers can then explore how these emotion displays are perceived by others, which conveys information about the expressiveness or communication of an emotion. There have been many studies of emotion recognition among autistic people (Harms, Martin, & Wallace, 2010). Some studies have identified that autistic people do not perform as well as TD people in labeling emotions (Baron-Cohen, Wheelwright, & Jolliffe, 1997; Celani, Battacchi, & Arcidiacono, 1999; Hobson, 1986; Tantam, Monaghan, Nicholson, & Stirling, 1989), whereas others report that there is no difference between the two groups (Prior, Dahlstrom, & Squires, 1990; Robel et al., 2004; Tanaka et al., 2012). In contrast, there have been comparatively few studies on the identification or interpretation of emotional expressions produced by autistic people (Brewer et al., 2015; Faso, Sasson, & Pinkham, 2015; Langdell, 1981; Macdonald et al., 1989; Volker, Lopata, Smith, & Thomeer, 2009). In the few studies that have been conducted, emotions have been elicited by asking the participant to pose for a certain emotion, and findings suggest that autistic people demonstrate a reduced ability to express emotions *via* their facial expressions compared to their TD peers (Yirmiya, Kasari, Sigman, & Mundy, 1989). It remains unknown, however, to what degree facial expressions of emotion and arousal among autistic people are present during naturalistic situations, and to what extent physiological arousal, facial expressions, and self-perception of anxiety are related in autism. Here, the authors present two studies that were designed to address these questions.

In Study 1, EDA, still video images of facial expressions, and self-report of anxiety were collected from autistic adolescents at each of 14 social skills intervention sessions offered by PEERS^®^ (Laugeson & Frankel, 2010). The goal of this study was to examine linkages amongst arousal and anxiety domains, rather than the effect of intervention on the domains themselves. Thus, the social skills intervention was primarily utilized to provide a naturalistic, ecologically-valid setting in which data could be collected repeatedly over a window of time.

Physiological arousal and the still facial expression images during intervention sessions were time-linked, such that facial expression still images from the highest and lowest physiological arousal points were extracted for further analysis. These facial expressions were then analyzed using machine learning algorithms in order to understand whether these algorithms could use facial expression features to predict which images were high arousal and which were low arousal. Further analyses examined the relations between physiological arousal and self-report of anxiety.

Given the results of Study 1, Study 2 was designed to address a competing hypothesis; instead of using machine learning algorithms, TD undergraduate students were asked to classify the still images of the autistic youth as high or low arousal state.

Results from Study 1 indicated that the adolescents experienced physiological arousal during intervention sessions (“Have-it”), and they were self-aware of it (“Know-it”), but the arousal state was not necessarily shown through their facial expressions (don’t “Show-it”). Classification results from Study 2 showed that just like the machine learning algorithms, humans were also not able to see the difference in arousal state from facial images of autistic adolescents.

## Methods

### Study 1

In Study 1, EDA, still video images of facial expressions, and self-report of anxiety were collected from autistic adolescents at each of 14 social skills intervention sessions. The goal of this study was to examine linkages amongst arousal and anxiety domains, rather than the effect of intervention on the domains themselves. Thus, the social skills intervention was primarily utilized to provide a naturalistic, ecologically-valid setting in which data could be collected repeatedly over a window of time.

#### Participants

Thirteen autistic male adolescents between the ages of 11 and 16 years were recruited for this study over the span of one and half years. Since this study was part of a larger randomized controlled trial (RCT; registered with ClinicalTrials.gov, Identifier: NCT02680015) of the Program for the Education and Enrichment of Relational Skills (PEERS^®^; (Van Hecke et al., 2013)), the inclusion criteria were as specified by prior work (Schohl et al., 2014). Briefly, autistic adolescents were required to score at or above a Composite IQ score of 70 on the Kaufman Brief Intelligence Test, 2^nd^ Edition (Kaufman & Kaufman, 2005), demonstrate verbal ability, and score at or above autism spectrum categorization on the Autism Diagnostic Observation Schedule-Generic, Module 3 or 4, (Lord, Rutter, DiLavore, & Risi, 1999). The Institutional Review Board at Marquette University approved both the overall study and this sub-study. Informed consent and, when possible, assent of the adolescents was obtained for the larger RCT and participation, with separate consent and assent, was offered for this additional, optional study. Adolescents and their parents were reminded that they could withdraw from the arousal-related tasks of the current study at any time, without affecting their participation in the social skills intervention. Adolescents were not compensated for arousal-related tasks, but the intervention was provided at no charge. One adolescent sat with his head down for several sessions, hence his video data was not analyzable for facial expressions. The data for this adolescent was thus dropped from the study, leaving data from 12 male adolescents for analysis. The characteristics of the adolescents are summarized in Table 2.

**Table 2.**
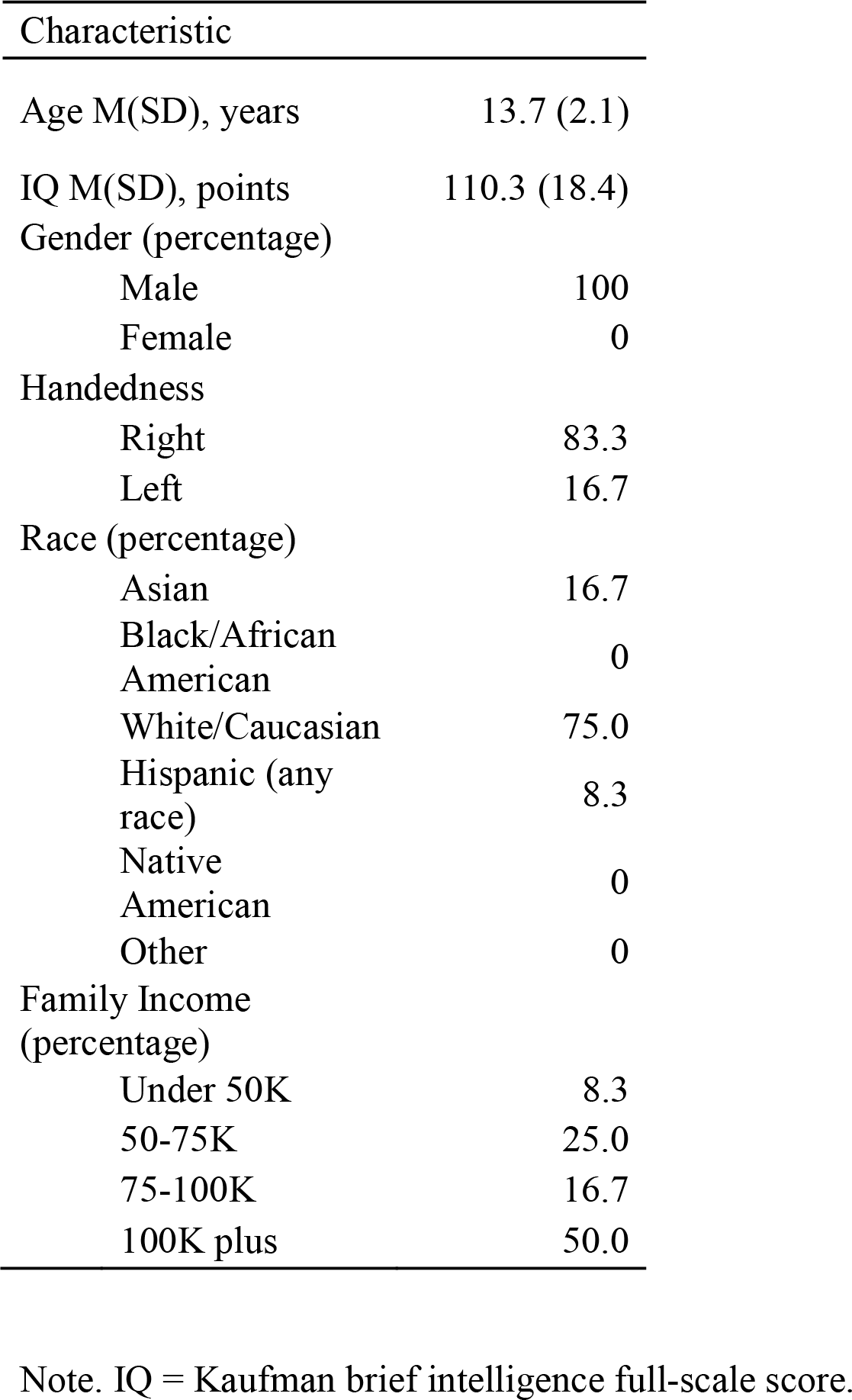
Participants’ Demographic Characteristics, Study 1

#### PEERS^®^ *I*ntervention *S*essions

Study 1 aimed to have a naturalistic environment for studying anxiety and physiological arousal among autistic adolescents, and PEERS^®^ sessions provided that. PEERS^®^ is a parent/caregiver-assisted social skills intervention for autistic adolescents (E. Laugeson, 2013; E. A. Laugeson & Frankel, 2011), which specifically targets skills pertaining to making and keeping friends, including starting and ending conversations, having get-togethers, and handling teasing and disagreements (E. A. Laugeson, Frankel, Mogil, & Dillon, 2009). Although there are variety of group-based social skills interventions for autistic youth (see Gates, Kang, & Lerner, 2017 for a recent review), PEERS^®^ is the only known intervention of its kind provided in this geographic region. Local societies offer support groups for families, but these do not include, to the authors’ knowledge, didactic (i.e., classroom-like) settings. The PEERS^®^ intervention, offered at the present institution, with a strong didactic component, provided the most viable option for data collection for this study.

The PEERS^®^ intervention consisted of 14 weekly 90-minute sessions with separate (concurrent) sessions for adolescents and parents/caregivers. Groups were led by trained graduate students (adolescent group leaders had at least a master’s degree in Psychology) under the supervision of a certified PEERS^®^ provider (the anchor author). For the first portion of the session, adolescents were seated around a table. Homework assignments for the week prior were reviewed, followed by a structured didactic lesson, along with role-plays of the target skill. The adolescents then participated in a behavioral rehearsal where they were coached in performing the target skill. Following rehearsal, adolescents and parents/caregivers were reunited and homework was assigned (for more details on the delivery of the intervention at this site, see (Schohl et al., 2014). Given potential issues with movement artifacts and off-camera time, EDA and video data were collected during the approximately 45-minute, structured, homework review and didactic portions of PEERS^®^, when adolescents were seated quietly. Although this program in itself has a strong evidence-base for use with autistic teenagers (E. A. Laugeson, Frankel, Gantman, Dillon, & Mogil, 2012), the uniqueness of this study lies in using PEERS^®^ with the aim of understanding anxiety and arousal among autistic adolescents in a naturalistic classroom-like setting over time.

### Measures

#### Physiological Arousal

Each PEERS^®^ cohort consisted of a maximum of ten adolescents. Two adolescents per cohort participated in this sub-study. Each adolescent participating in this sub-study was asked to wear a wristband called a Q-sensor, which is a wearable bio-sensor (Q-sensor: Affectiva, inc.). The adolescents wore the Q-sensor on their dominant wrist and the wristband collected continuous EDA data over the didactic portion (approximately 45 minutes) of the regular 90-minute PEERS^®^ session. As mentioned above, EDA is a measure of SNS activation and is measured in micro-siemens (μS). With the arousal of the SNS (fight/flight/freeze system), the sweat glands on the skin start producing sweat, which is followed by an increase in EDA. The data collected for each adolescent was saved as a file (see Figure 1a), which could be reviewed using the accompanied software. The software allows the user to export the file data to a spreadsheet (see Figure 1b), which can be used for mathematical data analysis.

**Fig. 1.**
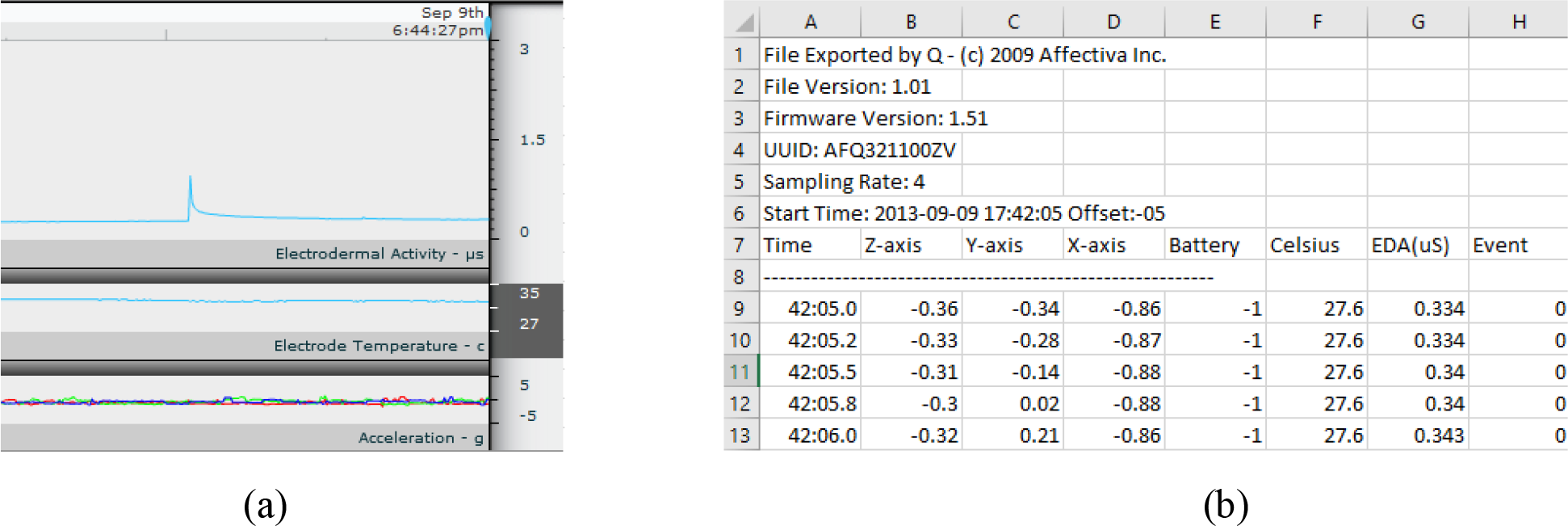
(a) Snippet of sample .eda file obtained from Q-sensor, (b) Equivalent .csv file imported using Q-sensor software

#### Facial Expressions

The autistic adolescents were recorded *via* two Sony (DCR-SR220) video cameras. One video camera was positioned approximately six feet in front of the adolescent so as to capture a full facial view. The second video camera was positioned to the left or right of the adolescent, at the rear of the room, and captured the overall events happening in the room, in order to provide contextual information. Just like the EDA data, video data from the cameras were recorded for all 14 sessions of the intervention. The video and EDA data were time-linked, so that the EDA events could be associated with specific points in time on the video.

#### Self-report of Anxiety

Autistic adolescents completed the short form of the Spielberg State-Trait Anxiety Inventory (STAI: (Marteau & Bekker, 1992) twice in each session: at the start and at the end of the didactic portion. This six-item form is a self-rating scale and has been validated for providing similar scores to those obtained using the full 20-item STAI (Marteau & Bekker, 1992). The six items include both anxiety-present (e.g., ‘I am tense’) and anxiety-absent (e.g., ‘I am relaxed’) items, and are rated on a four-point Likert scale, proving a total score. The measure has satisfactory reliability and is useful in cases of time constraint (Marteau & Bekker, 1992). Internal consistency for the total score was found to be acceptable (α = 0.78) in the present study.

### Procedure

#### EDA Data Analysis

The EDA data analysis consisted of calculating several parameters: maximum EDA, minimum EDA, mean EDA, area under the curve, standard deviation, and peaks per minute. These parameters were tabulated for each session and for each adolescent. Out of these parameters, maximum EDA was most useful in building the image dataset (see below). The additional parameters were also used in the regression analysis.

#### Facial Expression Analysis

The first step of facial expression analysis consisted of building a dataset of still facial expression images. These images corresponded to the exact moment in time when adolescents showed the highest level of EDA and the lowest level of EDA for a session. Specifically, for each adolescent, the time corresponding to the maximum EDA value in each session was calculated. Then, the facial image of the adolescent at that particular time in the video recording was captured (see Figure 2). This image was then labeled as the high EDA (“HiEDA”) image. When more than one instance of maximum EDA was found, preference was given to the instance with a higher clarity facial image. Similarly, a period of relatively low EDA was chosen, and a corresponding image was captured and labeled as the low EDA (“LoEDA”) image. This process was completed for all 12 adolescents, resulting in a total of 126 HiEDA and 126 LoEDA images. The total number of images for each adolescent differed, in that the number of PEERS^®^ sessions they attended varied due to absences. The number of images per adolescent ranged from 12 to 28, with an average of 21 images per adolescent.

**Fig. 2.**
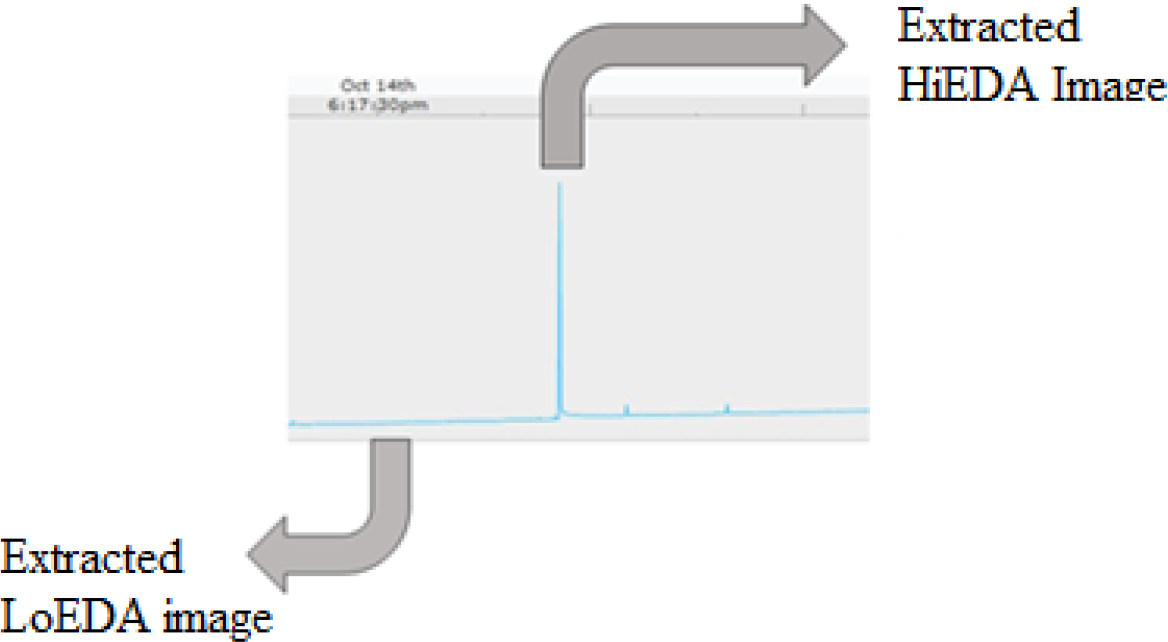
Preparing the Image Database - extracted still images at lowest and highest EDA. The facial images shown here have been intentionally turned into low resolution images to maintain adolescent’s anonymity.

The Eigenface method (Turk & Pentland, 1991) was used for extraction of facial expression information from each image in the image dataset. This method has predominantly been utilized as a face recognition method, however, in combination with classification techniques, it has also been used for facial expression recognition (Huang & Lin, 2008; Lau, 2010; Shinohara & Otsu, 2004). This method is based on Principal Component Analysis. Here, the facial images in the dataset were represented as vectors. Eigenvectors and eigenvalues were calculated from the covariance matrix and the set of eigenvectors was referred to as “eigenfaces.” Any facial image of a human can be written as a linear combination of these eigenfaces. The weights (w_1_, w_2_, w_3_, …) describe the contribution of each eigenface to a particular image. These weights were the projection of a particular image onto the face space and were calculated for all of the training dataset images and the test dataset images. The classification methods were then applied onto these weights for further understanding of facial images.

#### Image Clustering

The *k*-means clustering method (MacQueen, 1967) was applied to the image database to partition image data into groups. The *k*-means clustering method is a simple and widely-used clustering technique, where *k* is the number of desired clusters. Here, *k* was set to various values to assess the clustering tendency of the facial expression images. First, the value of *k* was set to the number of adolescents to check whether the images from the same adolescent would be clustered together or not. The value of *k* was also set to two to check if the images would be clustered according to the arousal level (i.e., HiEDA or LoEDA images). Because the PEERS^®^ intervention is comprised of 14 sessions where each session focuses on a different topic, the value of *k* was set to 14 to check if the images would be clustered by session. Finally, the optimal value for *k* was computed using the elbow method (Sugar, Lenert, & Olshen, 1999) and the analyzed the clustering results for the optimal *k* value.

#### Image Classification

In order to assess whether facial expression images could be classified into HiEDA and LoEDA statutes, supervised learning methods were employed to build image classifiers. In supervised learning, a labeled training dataset is used to build a classifier whose accuracy is computed using the held-out test dataset. Here, the image dataset was divided into training and test datasets. The training dataset was used to teach the classifier while the test dataset was used to evaluate the classifier. The algorithms used for classifying images into HiEDA or LoEDA were: *k-*Nearest Neighbor (*k*NN) algorithm, Support Vector Machine (SVM), and deep learning based Convolutional Neural Networks (CNN or ConvNets).

The *k*NN algorithm has been recognized as one of the simplest classification techniques in machine learning (Altman, 1992). Each of the samples in the training dataset has a class label. An unlabeled sample from the test dataset is assigned to the class by the majority of class votes from the *k* nearest neighbors. Here, *k* was set to three (i.e., an unlabeled sample was assigned to the class of majority of its three nearest neighbors from the training samples). The distance metrics used to find the nearest neighbors were ‘Euclidean distance’ and ‘cosine/angular distance.’

Support Vector Machines has been deemed one of the best techniques for binary classifications (Cortes & Vapnik, 1995). In the present study, the data had to be classified into two classes (HiEDA or LoEDA images), hence SVM technique was an appropriate choice. In SVM, the samples in the training dataset are divided into two classes by a hyperplane. Here, two different functions for this hyperplane – linear and quadratic – were used.

Deep learning is based on representation learning that allows a machine to automatically discover the features needed to classify the raw data (LeCun, Bengio, & Hinton, 2015). Deep learning based CNNs are particularly useful in image classification (Krizhevsky, Sutskever, & Hinton, 2012; LeCun, Bengio, & Hinton, 2015) and have been successfully used in facial expression recognition (Matsugu, Mori, Mitari, & Kaneda, 2003; Byeon & Kwak, 2014). Here, Keras – a deep learning library in Python (Chollet, 2017) – was used to build a CNN-based image classifier. The raw image files were used instead of the “eigenface” data. To avoid overfitting (i.e., the classifier achieves high accuracy on the training data, but low accuracy on the test data), the classifier complexity parameter, called epoch, was set to an optimal value. To determine the optimal epoch value, classifier accuracy on the training data was measured for different epoch values and the elbow method was used to pick the optimal epoch value. The optimal epoch value for the facial image dataset was five.

### Study 2

Study 2 was an extension of Study 1. In the first study, the facial images of the autistic adolescents were classified into high and low arousal images by machine learning algorithms. The average accuracy obtained in that case was slightly better than chance level. Thus, Study 2 explored whether humans were better at assessing arousal state than computers.

#### Participants

An IRB-approved study was conducted wherein 43 undergraduate students, aged ≥ 18, were recruited from Marquette University. The undergraduate students were consented for their participation in the study and received credit in psychology courses for their participation. The undergraduate students’ characteristics are shown in Table 3.

**Table 3.**
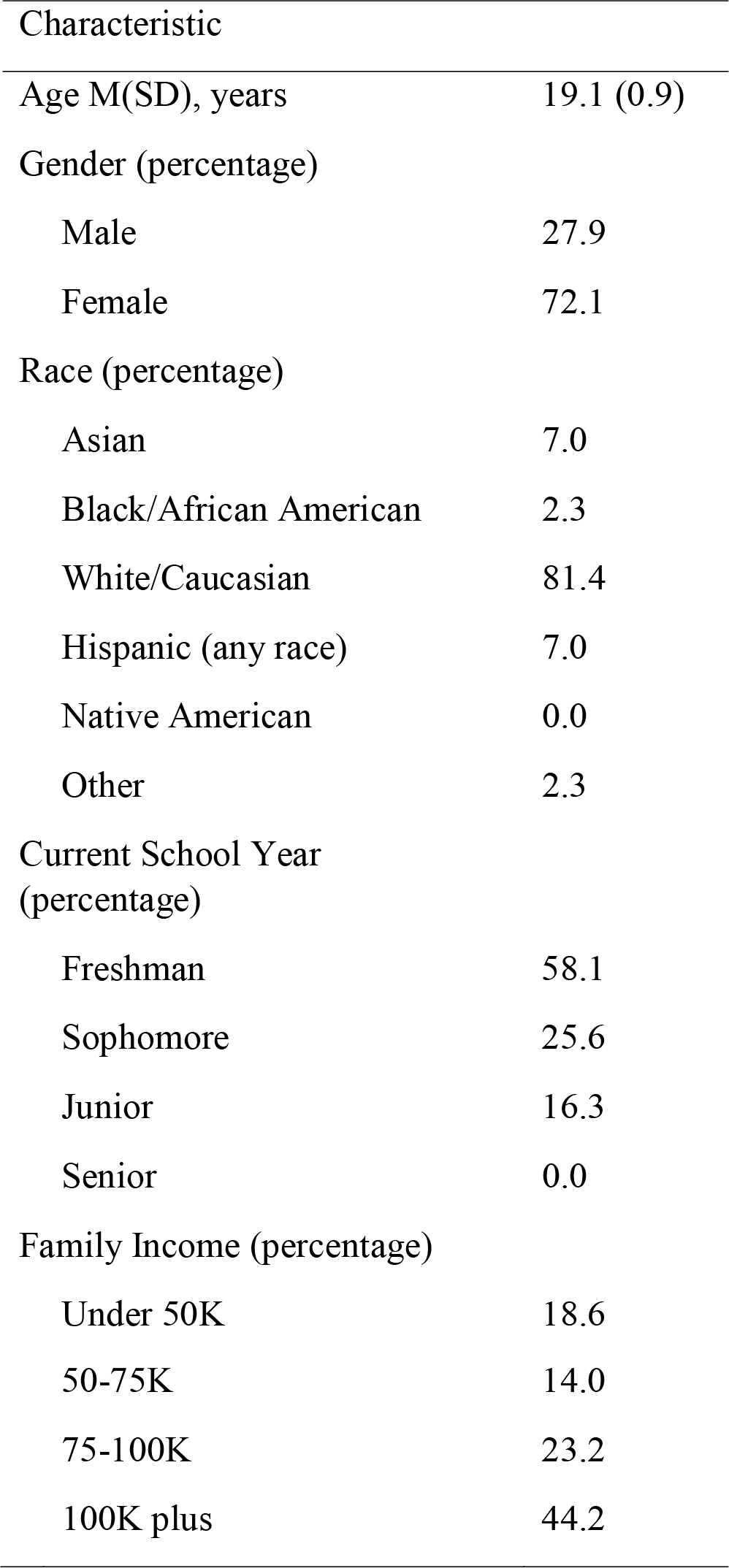
Participants’ Demographic Characteristics, Study 2

The autistic adolescents, whose images were to be shown to the undergraduate students, were separately consented for Study 2. That is, they and their parents/caregivers, were informed of the study and asked if they agreed to allow their/their child’s facial images to be shown to undergraduate students. Two out of 12 autistic adolescents did not consent for this study; hence their images were not included in Study 2. This reduced the image dataset from 252 images to 228 images.

#### Stimuli

The preparation of the image dataset for Study 2 was identical to Study 1. The image dataset was updated by removing images of autistic adolescents who did not consent for this study. The images from the updated image dataset (HiEDA and LoEDA images) were used in a randomized order.

#### Measures

To determine if humans were better at identifying the arousal of a person from their facial expressions, an online questionnaire was developed using Google Forms. This online questionnaire consisted of five items for undergraduate students’ demographic information, followed by the instructions related to the study. The undergraduate students were then directed to the Google Forms pages containing images from the image dataset. These images were to be rated on high/low arousal and positive/negative valence. Although data about ‘valence’ information of the facial image were collected, these data were not analyzed in the current study; only the arousal data are presented here.

#### Procedure

Data for this study were collected over four one-hour sessions, wherein 10-12 undergraduate students completed a session at a time. These sessions were conducted in a university computer lab, so as to accommodate a group of undergraduate students in each session. The undergraduate students received informed consented upon their arrival and were given a unique code to be entered in the online survey. Each undergraduate student was seated in a chair with a web-enabled IBM desktop in front of them for completing the online survey. A PowerPoint slide containing a brief explanation and examples of high/low arousal and positive/negative valence was projected onto a large screen at the front of the room for the duration of each session, such that it was easily visible to all of the undergraduate students and could be referred to in case of need. The undergraduate students were provided with a weblink to the online survey (Google Forms) containing images from the image dataset. For each image, undergraduate students had the options of choosing high vs. low arousal and positive vs. negative valence. The process was carried out for all of the images in the dataset, with the resulting undergraduate students’ responses saved in the underlying dataset for Google Forms.

## Results

### Study 1

#### EDA data analysis

Different EDA-based parameters (maximum EDA, minimum EDA, mean EDA, area under the curve, standard deviation, and peaks per minute) were observed for each adolescent during each session of the intervention. A wide variability was seen across the adolescents. As such, it could be that the content of each intervention session impacted the EDA of each adolescent differently. Session content may help to confirm this – some sessions focus on more challenging content, such as how to handle teasing or bullying, as opposed to potentially less intense topics like starting a conversation (Laugeson & Frankel, 2010). Maximum EDA for all 12 adolescents ranged from 0.014 to 11.37 μS (*M* = 1.962 μS, *SD* = 2.398).

#### *Image Clustering* using *k*-means Technique

The *k*-means clustering technique was used to cluster the image dataset. In the *k-means* clustering, *k* is set as the desired number of clusters. For experimental purposes, different values of *k* were chosen, but the most logical ones with respect to the dataset of the study were: *k* = 2, *k* = 12, and *k* = 14. This logic was based on the expectation that the image dataset could be clustered according to: 1) HiEDA and LoEDA images (*k* = 2), 2) the different faces in the dataset (i.e., the number of adolescents, *k* = 12), or 3) images from different sessions of the intervention (*k* = 14). Figure 3 shows the plots of *k*-means clustering results for *k* = 2, *k* = 12, and *k* = 14. It was observed that the clustering algorithm did not find any arousal level-, adolescent-, or session-based grouping among the images. For instance, in Figure 3b, each cluster had images belonging to several adolescents. Similar results were observed for two clusters (i.e., *k* = 2; Figure 3a) and fourteen clusters (i.e., *k* = 14; Figure 3c). This suggests that the variation in the images made recognition difficult for the clustering algorithms.

**Fig. 3.**
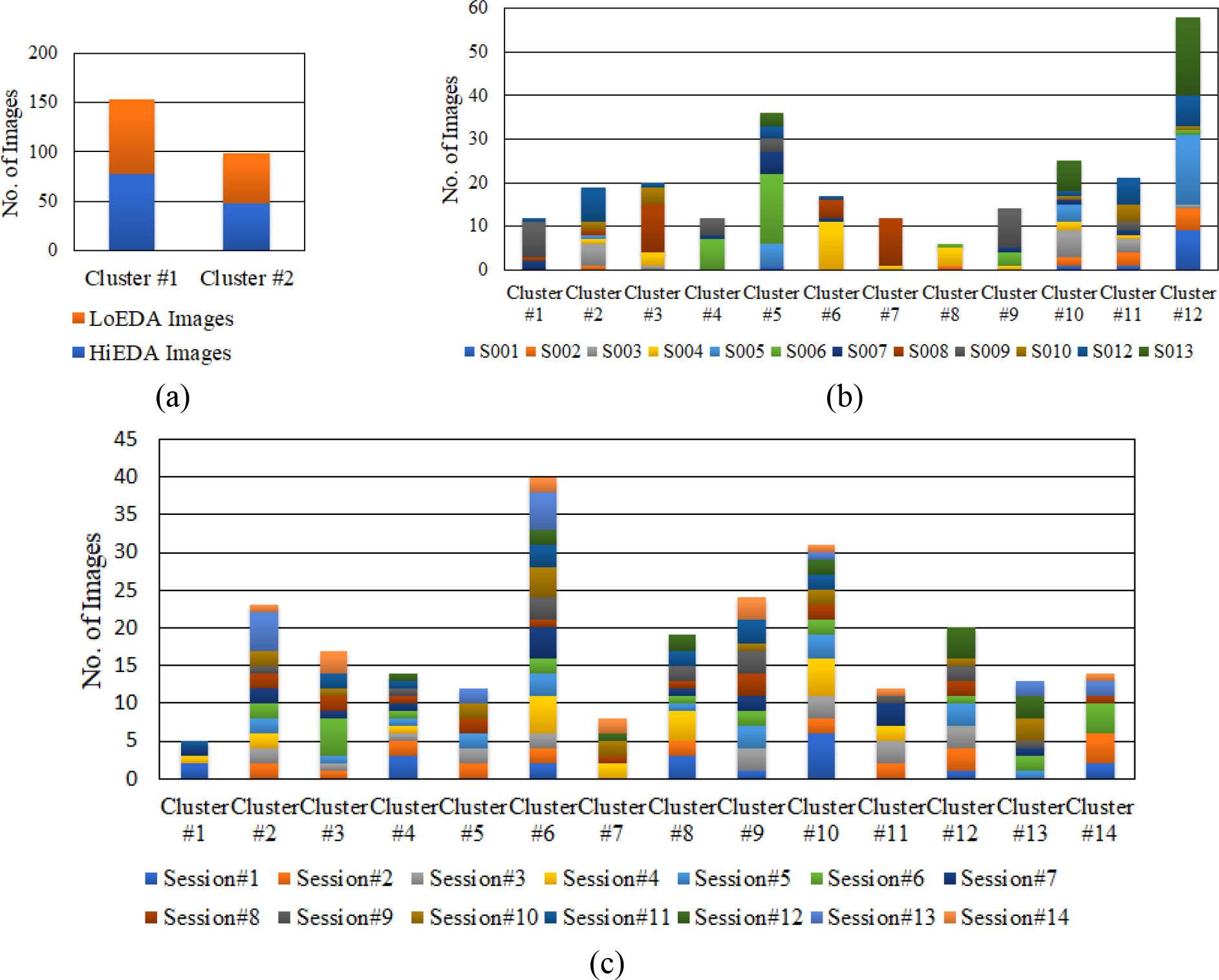
The k-means Clustering of Image Database for (a) *k* = 2, (b) *k* = 12, and (c) *k* = 14. The method was unable to identify discrete clusters of HiEDA/LoEDA images, individual adolescents’ images, and images from each session, respectively.

The optimal number of clusters to cluster the image data using *k*-means clustering, i.e., the optimal value for *k*, was found using the elbow method (Sugar, Lenert, & Olshen, 1999). Within-cluster sum of squared errors with respect to different *k* values were computed (Figure 4). The location of a bend (looks like an elbow) in the plot indicates the optimal value for *k*. In this case, the optimal *k*-value was three.

**Fig. 4.**
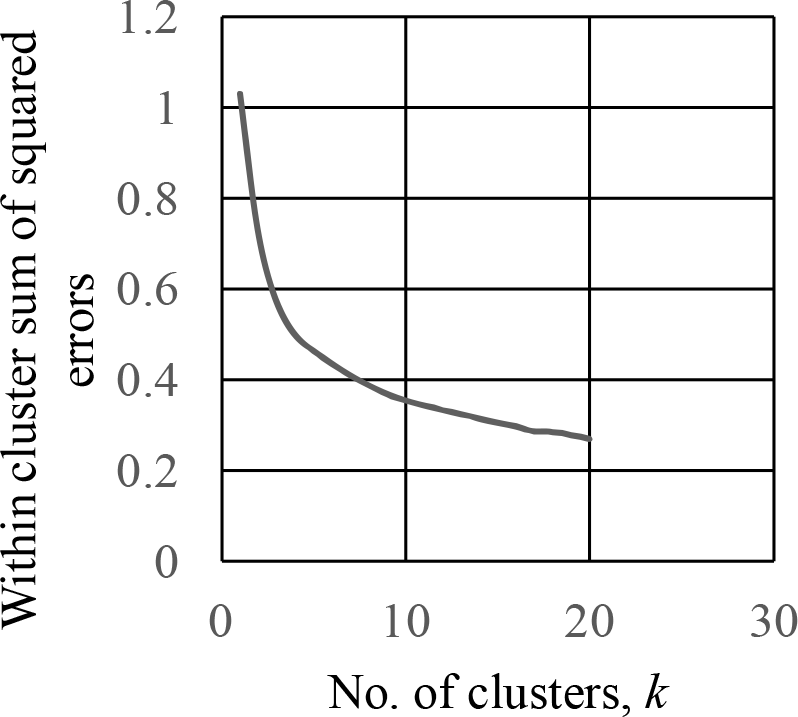
Finding the optimal value of *k* using the Elbow Method.

The *k*-means clustering technique was then applied for *k* = 3. It was observed that, for *k* = 3, there were no EDA- or session-based grouping (Figure 5a and Figure 5b). When the images in each cluster were labeled according to the adolescent they belonged to, however, it was observed that the images of some adolescents were grouped together in the same cluster (Figure 5c). For example, S005 had 26 images in the image database, 24 of them were in one cluster, and the remaining two were in another cluster. Similarly, S013 had all 28 images in one cluster. There were few adolescents, however, whose images were either (roughly) split into two clusters (e.g., S003 and S006) or all the three clusters (e.g., S010). Overall, the *k*-means clustering technique with three clusters seemed to recognize the facial features of different adolescents at a high-level.

**Fig. 5.**
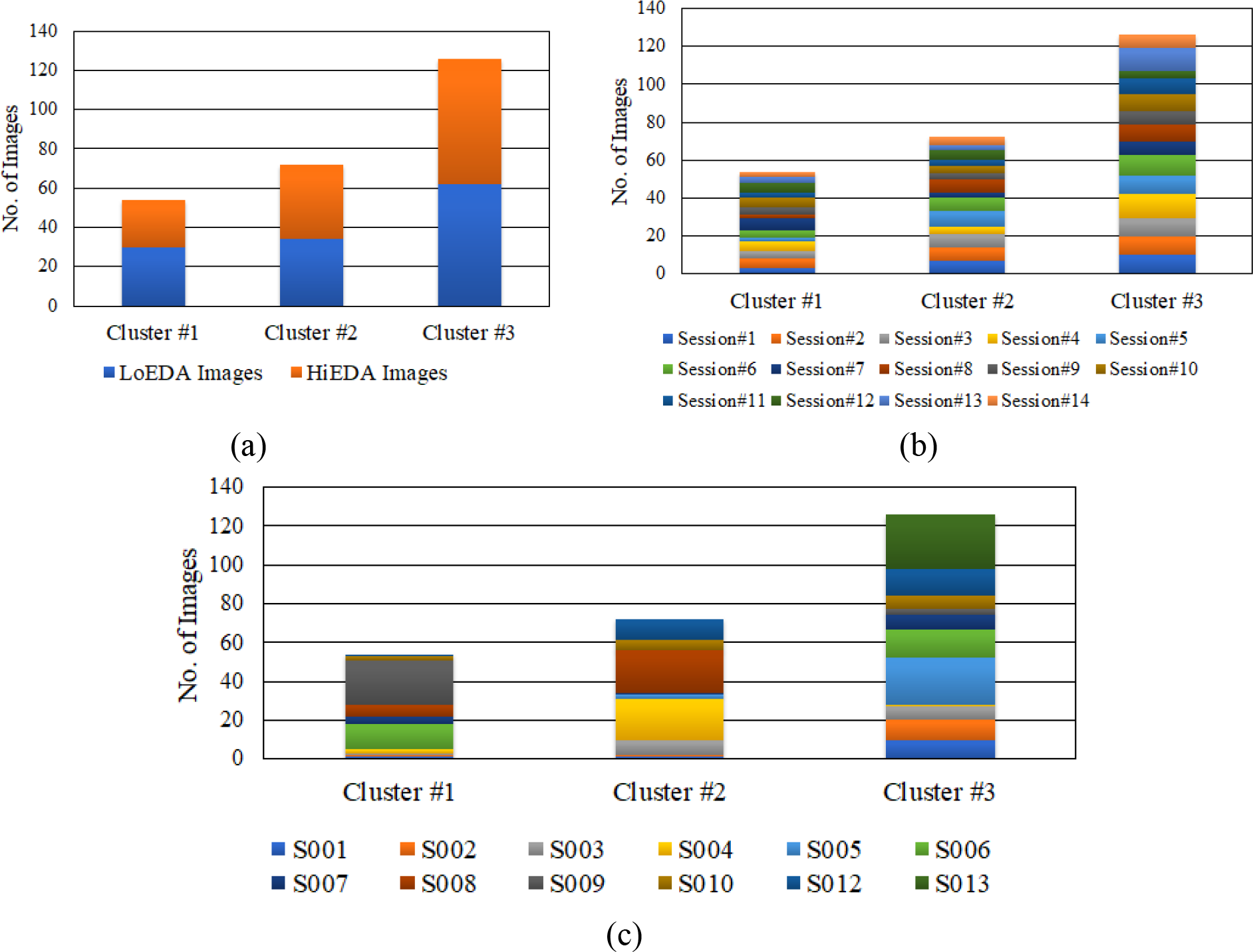
The k-means Clustering of Image Database (for *k=3*). The method was unable to identify discrete clusters of (a) HiEDA/LoEDA images, and (b) images from each session, respectively. The method could identify discrete clusters of individual adolescents’ images, (c) based on subject numbers (adolescents).

#### Image Classification using Leave-One-Subject-Out Cross-Validation

In order to assess whether image data could be classified into HiEDA and LoEDA statutes, supervised learning methods were applied to build image classifiers. In supervised learning, a classifer is trained using a subset of image dataset and tested to classify the held-out images. To check the performance of the classifier, leave-one-subject-out cross-validation approach was used. For this cross-validation approach, images of one adolescent were held out as the test images and images of all the remaining adolescents were used as the training images. The process was repeated until each adolescent’s image dataset was used as the test image. The classification was done using *k*-Nearest Neighbor (*k*NN), Support Vector Machine (SVM), and Convolutional Neural Networks (CNN) techniques (see Image Classification in Methods). The average accuracy of classification was then calculated (see Figure 6) and found to be 57.3%, slightly above the random chance level. Out of three techniques mentioned, the deep learning-based CNN approach gave the best average accuracy of 60.3%. These results indicate that the machine-learning-based methods found no strong relation between facial expression and arousal state of autistic adolescents.

**Fig. 6.**
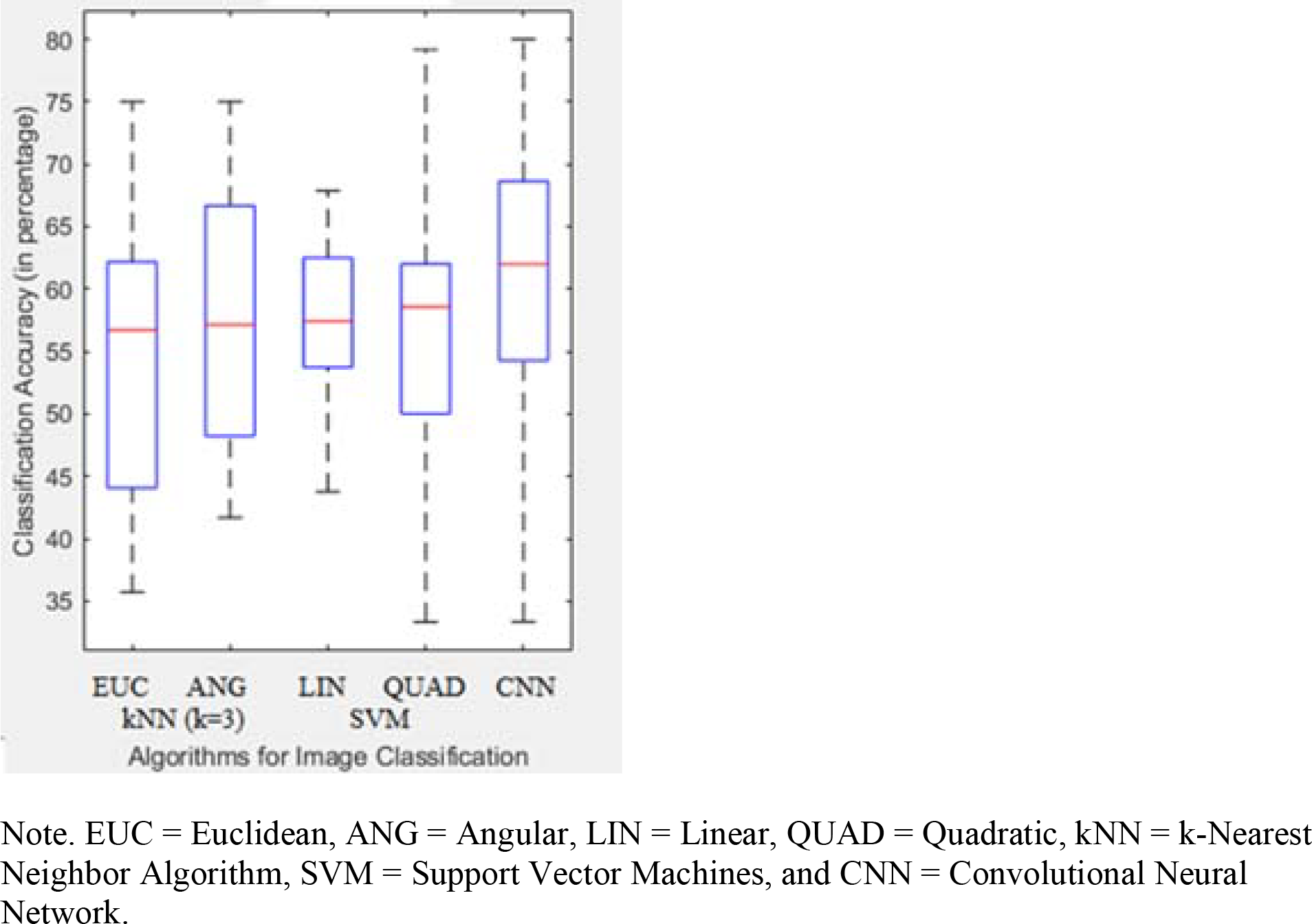
Classification accuracies of high and low EDA images, based on facial expressions, using Cross-validation method hovered between 50-70% accuracy, slightly better than chance.

#### Regression Analysis

A multiple linear regression analysis was performed to predict the average STAI total score (dependent variable) of each adolescent during each session from EDA parameters (mean EDA, maximum EDA, minimum EDA, area under curve, peaks per minute (ppm), and mean temperature). Session number was also included as an independent variable. This analysis indicated that session number and minimum EDA values had significant relations to average STAI total score, *F*(10,122) = 4.3, *p* < 0.05, *β*_session#_ = −0.2, *β*_minEDA_ = 2.3. These results demonstrate that the autistic adolescents were self-aware of their arousal states, on average. Results also suggest that the session number, that is, the topic for the intervention session, or the day in general, had an impact on the arousal state of the adolescents.

### Study 2

The 228 images in the Study 2 image dataset were rated on high/low arousal level by 43 undergraduate students. The reliability of agreement between undergraduates when assigning a high or low arousal category to these 228 images was assessed using Fleiss’ kappa coefficient (Fleiss, 1971). Kappa was found to be 0.23 on a scale of −1.00 to 1.00, indicating a low level of agreement between the raters. The average classification accuracy for the image dataset was found to be 51.1%, ranging from 44.3% to 57.9%. This classification accuracy was poorer than that obtained from the machine learning algorithms (57.3%), indicating that humans were no better than the computer algorithms at judging arousal state from the facial expressions of the autistic adolescents.

## Discussion

In Study 1, the first goal was to elicit physiological arousal among autistic adolescents in a natural way. The EDA data collected using the Q-sensor wristbands were helpful in identifying the periods of highest and lowest arousal during each session of the intervention.

The second goal of Study 1 was to see if this arousal was evident from the facial expressions of autistic adolescents as evaluated by machine learning algorithms. A *k*-means clustering algorithm was employed to cluster the facial image data and several supervised learning algorithms (*k*NN, SVM, and CNN with different parameters) were employed to classify the facial image data. These algorithms were able to predict the EDA arousal states (high and low) with slightly better than chance accuracy (57.3%). Deep learning-based CNN gave the best average accuracy compared to *k*NN and SVM techniques. The clustering technique using *k*-means was not able to clearly cluster the image data according to arousal states or the autistic adolescents. This result led to the conclusion that the linkage between facial expressions of autistic adolescents and their physiology was weak. When the change in different facial expressions was close to non-existent, the machine learning algorithms were not able to spot the difference in high and low arousal states with clarity. This could be the reason behind low classification accuracy of image dataset.

The third goal of Study 1 was to see if the autistic adolescents were self-aware of their anxiety. For this, a regression analysis between the minimum EDA and average anxiety score showed a significant relation. We also observed a significant relation between session number and average anxiety score. The overall results of Study 1 led to the conclusion that there were instances when physiological arousal was experienced by the adolescents (“Have-it”), of which they were self-aware (“Know-it”), but which they did not necessarily express through their facial expressions (don’t “Show-it”).

The goal of Study 2 was to compare the machine algorithms’ accuracy to human accuracy in identifying arousal from facial images of the autistic adolescents. A comparison of the average accuracies of classification obtained from the machine learning algorithms (57.3%) and humans (51.1%) led to the conclusion that humans were no better at predicting arousal levels from the facial expressions than the machine learning algorithms. Again, this finding could be because of the lack of change in facial expression from one arousal state to another (high to low, or vice-versa) of the autistic adolescents. Moreover, the inter-rater reliability, found using Fleiss’ kappa coefficient, indicated a poor level of agreement among the undergraduate students. This finding indicated that the undergraduate students were not consistent among themselves in rating an image as high or low arousal.

One of the limitations of Study 1 was its small sample size. The design of this study was such that with more adolescents the need of more wristbands and video cameras emerges. This requirement poses financial as well as practical constraints on the experimental setting. Deep learning methods have shown excellent results with large training datasets (Krizhevsky, Sutskever, & Hinton, 2012; Russakovsky et al., 2015), for instance, containing 1.3 million images in a training set classified into 1000 classes. Thus, with more training data, deep learning-based methods might detect subtle relations between facial expressions and arousal levels more accurately.

Another limitation was the lack of a TD comparison group. One major challenge of having such a group would be creating the same environment for stimuli and data collection for the control group. Future studies should attempt to address this limitation.

Study 2 also lacked contextual information for the undergraduate students to utilize while rating the facial images on arousal level. Research has shown that context plays an important role in observers’ understanding of everyday emotional experiences (Hassin, Aviezer, & Bentin, 2013). If one tries to understand facial expressions (in this case, specific to arousal) from still images, one is very likely to miss important information available through the context. It might be helpful for participants to view the entire video segment and annotate the affective state of a person during the PEERS^®^ intervention. Moreover, this annotation, if done by a behavioral expert, could add a more reliable perspective to the research.

## Conclusion

There is a very limited literature examining the physiology of autistic adolescents, especially in connection with affective facial expressions. This study provides preliminary evidence pertaining to the relation between “have-it,” “know-it,” and “show-it” components of physiological arousal among autistic adolescents. Even with its limitations, this study provides a better understanding of the expression of arousal and anxiety in autism and provides a foundation for future research in this area, such as treatment effects on these domains. Further, the findings of this study are informative for clinical and school settings, in which the anxiety and arousal experienced by autistic youth may affect their functioning. It is important to consider that, although not readily shown on the face, autistic youth may experience a wide range of arousal and anxiety, and they may be self-aware of these states. For example, it is often reported that autistic children and adolescents have “sudden” incidents of behavioral dysregulation (i.e., “meltdowns”), where it is difficult for others to identify antecedents of these states. The results of the current study suggest that these instances may not be “sudden,” but rather not communicatively expressed in advance. If dysregulation is not ameliorated, it may be that autistic youth become more and more distressed, eventually reaching the point of “meltdown.” Utilizing non-invasive, nonverbal, physiological monitoring technology may assist autistic youth in communicating their increasing arousal earlier, which may pave the way for better regulation, and ultimately, better functioning across a variety of settings.

## Acknowledgements

The authors would like to thank the autistic adolescents and their families as well as Marquette undergraduates for their participation in our research. The authors also recognize the Marquette Autism Project team for their work on this project and in the lab. Finally, the authors would like to extend their gratitude to Elizabeth A. Laugeson, Psy.D. for her assistance in offering the PEERS^®^ interventions in Wisconsin. A portion of this project was presented as a poster presentation at the International Meeting for Autism Research in Salt Lake City, May 2015 under the title, “Physiological Monitoring During PEERS^®^: A ‘Culture-Free’ Method of Understanding Intervention Response.” This paper was originally prepared as the doctoral dissertation for the lead author: Niharika Jain, under the title, “Affective Computing in the Area of Autism.”

## Footnotes

* We use the term “autistic person” rather than “person/child/adolescent with autism” to respect a preference in the autism community for the use of identity first language. For more information, see https://autisticadvocacy.org/about-asan/identity-first-language/.

